# Genetic Dynamics of Mustang and Feral Horse Populations in the Western United States

**DOI:** 10.1101/2024.01.28.577652

**Authors:** E. Gus Cothran, Anas Khanshour, Stephan Funk, Eleanore Conant, Rytis Juras, Brian W. Davis

## Abstract

The history and population dynamics of feral horse and wild mustang population in the Western United States has led to diverse populations of disparate ancestry. These iconic populations are currently managed by the Bureau of Land Management (BLM) and their genetic history is of great interest for both management and conservation purposes. We examined population genetic parameters using 12 well established microsatellite loci in nearly 8,500 horses representing 235 populations sampled across more than 20 years. Samples were collected by BLM or by members of other management agencies from 10 states. Genetic variability and genetic resemblance to domestic horse breeds using multiple methods were estimated. A wide range of variation levels were observed across the populations. In general, within-population variability was slightly lower than what has been found in domestic horse breeds, but still retains diversity. As expected, levels of population variation correlated to census size. Several populations were sampled longitudinally with intervals between sampling of about 5 years. For these longitudinal samples, there was no trend towards an increase or decline in diversity, indicating consistent management practices. Relationships between populations and domestic breeds ranged from close association to one or two specific breeds to extreme divergence of the feral horses to all breeds examined. Reasons for divergence are mainly related to the founding of the population and subsequent demographic history. Overall, there was a slight tendency for geographically close feral populations to be more similar to each other than to more distant populations. The results of this study show the feral horse populations in the western US have a considerable variation, though management practices can strongly influence variability levels.

## Materials and Methods

We tested over 8,520 horses representing 235 populations. Hair follicles or blood samples were collected by BLM personnel or by members of other management agencies from 10 states (summary statistics of herds per state are shown in Table 1 and the location of feral herds in the US is shown in Figure 1). Average sample size per population was about 38 horses. Some populations were sampled two or more times over the covered by this study which is over 20 years. Total DNA was extracted from hair follicle samples using the PUREGENE® DNA purification kit following the manufacturer’s protocol. Microsatellite genotyping was performed using an ABI PRISM 3730 *xl* (Applied Biosystems, Foster City, CA, USA) following previously described methods (Juras, Cothran and Klimas, 2003; Khanshour*, et al.*, 2013).

**Figure 1.**
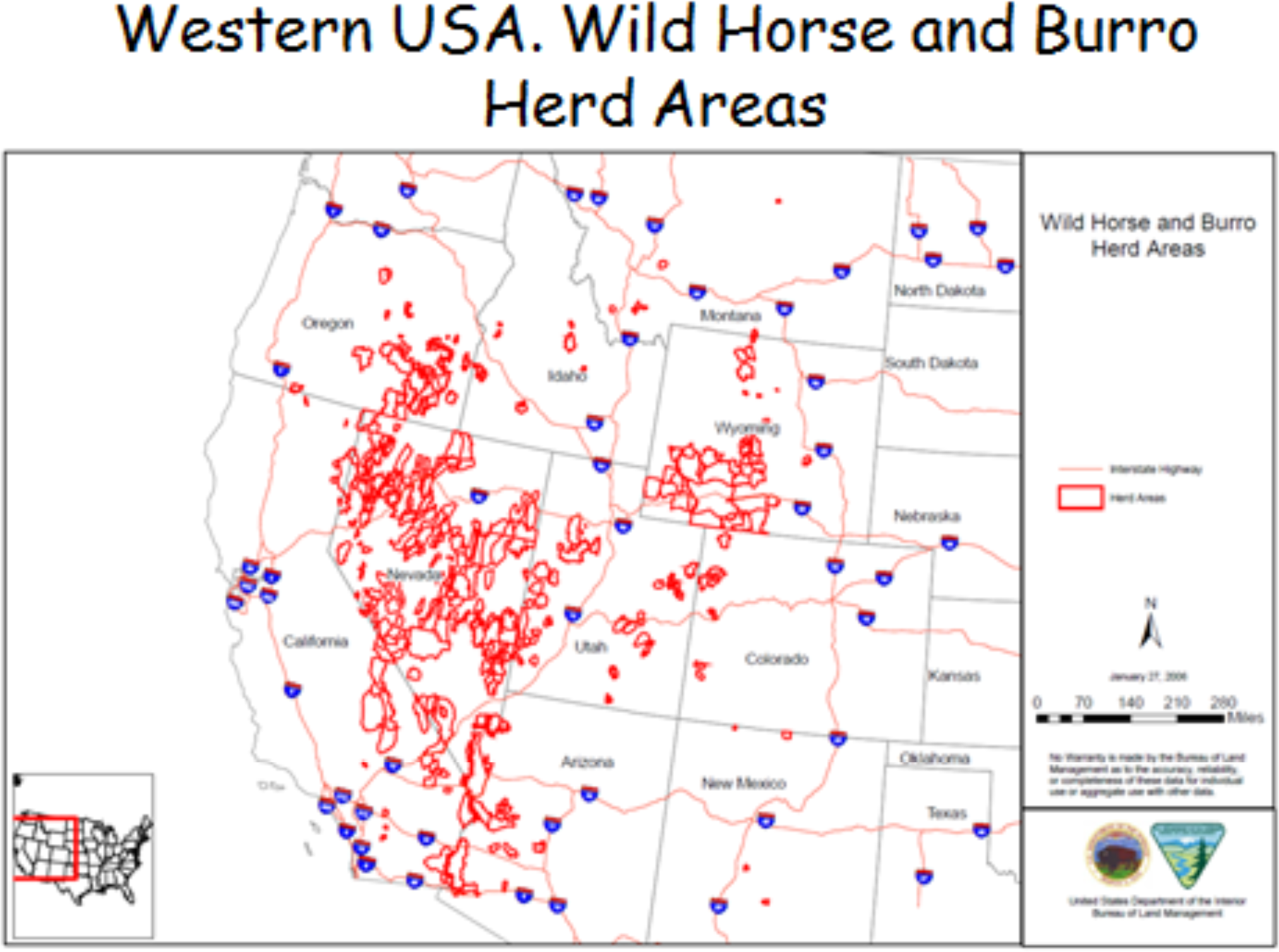
Location of herd management areas in the western US.

We examine 12 autosomal microsatellite markers distributed across the horse genome. We calculated genetic diversity indices for each herd using GENEALEX 6 (Peakall and Smouse, 2012). These included observed heterozygosity (Ho), expected heterozygosity (He), estimated inbreeding level (Fis), Nei’s unbiased heterozygosity (Hub), effective number of alleles (ENA), total number of alleles (NA), Mean number of alleles per locus (MNA), number of alleles at less than 5% frequency (termed as rare alleles here; RA), percentage of rare alleles (RARE) and mean rare alleles per locus (MRA).

For the following analyses 152 herds were used. For herds that were sampled multiple times only a single sampling period was used and only BLM herds were included except for two Forest Service herds from New Mexico. The genetic distance among populations was investigated by the Majority-rule consensus of Restricted Maximum Likelihood (RML) trees. We calculated the chord distances generated from 1000 bootstrapped allele frequency datasets using the CONTML and CONSENSE procedures in the PHYLIP 3.69 package (Felsenstein, 1989) http://evolution.genetics.washington.edu/phylip.html.

Trees were visualized by MEGA4 (Tamura*, et al.*, 2007). The Przewalski Horse was used as an out-group. Genetic distance was compared to geographic distance using the Mantel matrix correlation test in the SAS statistical package. We used a geographic information system (GIS; ArcMap 10, ESRI, Redlands, California) to determine the pairwise geographic distance between sampled administrative units (herd management areas, herd areas, and wild horse territories). Edge-to-edge distance was used for this measure, because it was assumed that horses could have used any part of that unit in which they were caught. Contiguous units had distance values of zero. For all other units, the pairwise distance used was the minimum distance between the two units’ boundaries. The genetic distance used was Nei’s modified distance *Da* (Nei, 1978).

To represent the structure and differentiation among herds, pairwise Fst values were calculated using Arlequin 3.5.1.3 (Excoffier and Lischer, 2010). We also ran the Principal Coordinate Analysis (PCoA) based upon the dissimilarity matrix of the Chord Distance using DARwin 5.0 (http://darwin.cirad.fr/darwin). Furthermore, we tested the evidence of immigration into most herds by comparing the fit of each individual to a panel of reference breeds which included the population being examined using the program WHICHRUN (Banks and Eichert, 2000). Only selected herds were included in this analysis.

STRUCTURE 2.4.4 (PRITCHARD, STEPHENS AND DONNELLY, 2000) was used to infer genetic structure for 160 herds (the additional herds were mainly non-BLM herds not included in other analyses) assuming K=1 to K=12 ancestral populations. Analyses utilized correlated allele frequencies, applied information on the sample origin (with locprior) and allowed admixture accounting for recent gene flow between populations (Funk, *et al.*, 2020). A total of 1,500,000 MCMC including 1,000,000 burn-in iterations was selected as the setting did not produce results differing from larger MCMC iterations in the data set of all breeds (Funk, *et al.*, 2020). MCMC runs were repeated 20 times. For each run, posterior probabilities of the data were checked visually for convergence over increasing numbers of iterations (10). Visual presentation of the clusteŕs estimated membership coefficients for individuals and populations utilized DISTRUCT 1.1 (ROSENBERG, 2004) for the highest posterior probability of the data at each K. We evaluated visually how the range of K=2 to K=12 ancestral populations can explain the data in relation to known natural history data from the populations rather than using a point estimate such as the Evanno et al. (Evanno, Regnaut and Goudet, 2005) ΔK estimate. Funk et al. (Funk, *et al.*, 2020) demonstrated with a large dataset of horses that such point estimates can be problematic lending support for forfeiting point estimates of an increasing number of studies including ones covering horses (Cortés, *et al.*, 2017) or sheep (Leroy, *et al.*, 2015). All STRUCTURE analyses were carried out on an SGI UV 2000 computing platform with 96 nodes, 192 cores and 1.5 teraflop.

FLOCK 3.1 (Duchesne and Turgeon, 2012) was applied using 20 re-allocations for the multilocus maximum likelihood procedure with 50 runs for each population cluster K and a LLOD threshold score of 0. The optimal number of clusters that explain the data was determined by ad hoc “stopping” rules (Duchesne and Turgeon, 2012). Clusters are hierarchically evaluated starting at K=2; K is stepwise increased until no stopping condition is found for four successive values of K. If no stopping condition is found at all, i.e. an “undecided stopping condition”, then this may be due to the absence of genetic structure or too low a genetic information content of the genotyped loci (Duchesne and Turgeon, 2012). For the horse microsatellite loci we used, stopping conditions were encountered in a different breed set (Funk, *et al.*, 2020), demonstrating that the information content of the marker set is sufficient: thus, a “undecided stopping condition” means in this context that no genetic structure could be detected in the current data set.

Modularity analysis was performed to understand the division of a network, in this case a series of interconnected horse populations, into modules or groups. Nodes, for our purposes are concentrations of individuals in habitat patches, within a module are connected with respect to their genetic network. This connectivity is the primary metric affecting modularity, as measured by within-module strength and participation coefficient, which measures the weight of a patch in connectivity maintainance within and among modules.

To accomplish this, we used the multilocus microsatellite data after filtration for presence of all 12 loci in all individuals, and consistent record of sampling latitude and longitude, resulting in 6070 samples. Code was modified based on the integrated analysis in Peterman *et al*. (Peterman, *et al.*, 2016), and relied on the methods of Dyer and Nason (Dyer and Nason, 2004). All calculations were performed using the gstudio and popgraph packages in R 4.0 (http://CRAN.R-project.org/package=popgraph). We constructed an incidence matrix of genetic covariance that identifies the most important connections between populations, which created the network that we subsequently evaluated for modularity. Modularity evaluation for our horse genetic network closely followed the methods of Fletcher *et al*.(Fletcher, *et al.*, 2013) and the equation described by Girvan and Newman (Girvan and Newman, 2002).

## Results and Discussion

Mean measures of variability for all populations and for domestic breeds are shown in Table 2 and the total data set giving all measures for each population in Table 1. Observed heterozygosity ranged from 0.497 to 0.815 which essentially spans the range of variation for domestic breeds. The same could be said for all variability measures. However, on average, the different estimators for genetic variation levels tend to be lower in the feral horse populations than in domestic breeds (Table 2). This likely due to small population size and founder effect. A variety of factors can influence genetic variability in these relatively small populations. In addition to variability measures we obtained estimates of the percentage of individuals within the population that do not fit the genetic profile of the population and may represent recent immigrants into the herd. Also, we obtained the Appropriate Management Level (AML) for each herd. This is the number of horses that the BLM calculates that the herd area can support and thus is an estimate of population size (actual census numbers are almost never available). The herd will increase in size above the AML between “gathers” (round-ups of horses which are mainly used to remove what are considered to be excess animals) but the population size is usually reduced to a number below the AML at the time of a gather.

**Table 2.** Mean genetic variability measures for feral and domestic horse populations.

| Type | Variable | N | Mean | Std Dev | Minimum | Maximum |
| --- | --- | --- | --- | --- | --- | --- |
| Feral Herds | SIZ | 229 | 37.84 | 29.31 | 4 | 209 |
|  | Ho | 229 | 0.722 | 0.059 | 0.492 | 0.867 |
|  | He | 229 | 0.718 | 0.059 | 0.448 | 0.801 |
|  | Fis | 229 | -0.008 | 0.071 | -0.284 | 0.382 |
|  | Hub | 229 | 0.733 | 0.055 | 0.462 | 0.808 |
|  | AE | 226 | 3.965 | 0.646 | 1.988 | 5.253 |
|  | NA | 229 | 74.948 | 13.233 | 31 | 97 |
|  | MNA | 229 | 6.246 | 1.103 | 2.583 | 8.083 |
|  | RA | 229 | 18.345 | 8.024 | 0 | 34 |
|  | RARE | 229 | 0.234 | 0.087 | 0 | 0.4 |
|  | MRA | 229 | 1.529 | 0.669 | 0 | 2.833 |
| Domestic Breeds | SIZ | 211 | 66.22 | 124.81 | 2 | 1195 |
|  | Ho | 211 | 0.725 | 0.073 | 0.264 | 0.879 |
|  | He | 211 | 0.722 | 0.066 | 0.213 | 0.812 |
|  | Fis | 211 | -0.007 | 0.079 | -0.551 | 0.286 |
|  | Hub | 211 | 0.737 | 0.063 | 0.232 | 0.829 |
|  | AE | 209 | 4.021 | 0.714 | 1.409 | 5.599 |
|  | NA | 211 | 79.588 | 16.365 | 20 | 119 |
|  | MNA | 211 | 6.632 | 1.364 | 1.667 | 9.917 |
|  | RA | 211 | 22.427 | 10.777 | 0 | 55 |
|  | RARE | 211 | 0.267 | 0.099 | 0 | 0.462 |
|  | MRA | 211 | 1.869 | 0.898 | 0 | 4.583 |

Variability was strongly associated with population size as measured by AML. All measures of variability were significantly (Figure 8) and positively associated with AML. Only Ho had a relatively low association with AML but even this was statistically significant at the p = 0.02 level. Relationships for each measure with AML are shown in Figure 8. Population sizes of the herds have fluctuated over time. When a herd was sampled on more than one occasion, levels of variation at the different time periods were compared (Table 3). In most cases no large change was evident, mainly because sampling periods spanned less than a generation interval (see below). However, some populations showed increases in variability over time (for example OR0014 in 2001 compared to 2011) while others showed decreases; (for example NV0226 2001 compared to 2012) and there was no consistent trend across all multiply sampled herds. Some herds that were sampled three times had variation levels that increased from the first time period to the second then decreased in the third (NV0103 and NV0222 for example). There also was a significant association of the estimate of the number of migrants in a population and genetic variation with variation increasing as the number of migrants increased (Figures 9).

**Table 3:** Genetic variability measures for herds tested at two or more time periods.

| **BNAM** | **Herd** | **YEAR** | **N** | **Ho** | **He** | **ENA** | **NA** | **MNA** | **%Rare** |
| --- | --- | --- | --- | --- | --- | --- | --- | --- | --- |
| CA0242 | TWIN PEAKS CA | 2001 | 48 | 0.780 | 0.798 | 5.254 | 95 | 7.917 | 0.274 |
| CA0242 | TWIN PEAKS CA | 2011 | 93 | 0.701 | 0.768 | 4.826 | 97 | 8.083 | 0.289 |
| CA0262 | BUCKHORN CA | 2003 | 28 | 0.798 | 0.764 | 4.524 | 80 | 6.667 | 0.225 |
| CA0262 | BUCKHORN CA | 2010 | 31 | 0.806 | 0.749 | 4.155 | 77 | 6.417 | 0.208 |
| CA0269 | CARTER RESERVOIR CA | 2003 | 26 | 0.670 | 0.650 | 3.108 | 57 | 4.750 | 0.088 |
| CA0269 | CARTER RESERVOIR CA | 2009 | 60 | 0.689 | 0.688 | 3.475 | 76 | 6.333 | 0.263 |
| CO0162 | WEST DOUGLAS CO | 2001 | 32 | 0.753 | 0.712 | 3.568 | 70 | 5.833 | 0.286 |
| CO0162 | WEST DOUGLAS CO | 2006 | 35 | 0.686 | 0.691 | 3.329 | 55 | 4.583 | 0.091 |
| CO0766 | LITTLE BOOKCLIFFS CO | 2002 | 29 | 0.745 | 0.721 | 4.125 | 75 | 6.250 | 0.213 |
| CO0766 | LITTLE BOOKCLIFFS CO | 2013 | 19 | 0.715 | 0.684 | 3.775 | 71 | 5.917 | 0.169 |
| ID0005 | CHALLIS ID | 2002 | 46 | 0.743 | 0.735 | 3.938 | 77 | 6.417 | 0.247 |
| ID0005 | CHALLIS ID | 2012 | 39 | 0.733 | 0.724 | 3.734 | 75 | 6.250 | 0.240 |
| ID0007 | HARDTRIGGER ID | 2003 | 25 | 0.699 | 0.749 | 4.286 | 79 | 6.583 | 0.190 |
| ID0007 | HARDTRIGGER ID | 2010 | 30 | 0.764 | 0.733 | 4.128 | 78 | 6.500 | 0.282 |
| MT0251 | PRYOR MOUNTAINS MT | 2001 | 209 | 0.715 | 0.693 | 3.917 | 90 | 7.500 | 0.367 |
| MT0251 | PRYOR MOUNTAINS MT | 2009 | 103 | 0.757 | 0.762 | 4.507 | 79 | 6.583 | 0.190 |
| MT0251 | PRYOR MOUNTAINS MT | 2012 | 45 | 0.720 | 0.728 | 4.008 | 75 | 6.250 | 0.200 |
| NM0001 | BORDO ATRAVESADO NM | 2011 | 27 | 0.787 | 0.749 | 4.338 | 73 | 6.083 | 0.164 |
| NM0001 | BORDO ATRAVESADO NM | 2012 | 30 | 0.742 | 0.726 | 4.014 | 70 | 5.833 | 0.200 |
| NV0102 | LITTLE HUMBOLT NV | 2002 | 17 | 0.779 | 0.728 | 4.158 | 72 | 6.000 | 0.153 |
| NV0102 | LITTLE HUMBOLT NV | 2010 | 23 | 0.743 | 0.742 | 4.312 | 79 | 6.583 | 0.266 |
| NV0103 | ROCK CREEK NV | 2002 | 25 | 0.687 | 0.700 | 3.758 | 67 | 5.583 | 0.194 |
| NV0103 | ROCK CREEK NV | 2010 | 28 | 0.738 | 0.745 | 4.123 | 72 | 6.000 | 0.153 |
| NV0103 | ROCK CREEK NV | 2016 | 30 | 0.717 | 0.729 | 3.926 | 76 | 6.333 | 0.289 |
| NV0107 | ANTELOPE VALLEY NV | 2002 | 48 | 0.673 | 0.746 | 4.144 | 90 | 7.500 | 0.367 |
| NV0107 | ANTELOPE VALLEY NV | 2011 | 28 | 0.765 | 0.756 | 4.305 | 81 | 6.750 | 0.235 |
| NV0108 | GOSHUTE NV | 2001 | 20 | 0.741 | 0.720 | 3.878 | 72 | 6.000 | 0.292 |
| NV0108 | GOSHUTE NV | 2011 | 28 | 0.765 | 0.748 | 4.300 | 78 | 6.500 | 0.205 |
| NV0220 | BUFFALO HILLS NV | 2002 | 16 | 0.668 | 0.679 | 3.440 | 68 | 5.667 | 0.206 |
| NV0220 | BUFFALO HILLS NV | 2009 | 51 | 0.724 | 0.729 | 4.029 | 84 | 7.000 | 0.286 |
| NV0221 | GRANITE RANGE NV | 2002 | 47 | 0.774 | 0.731 | 4.110 | 82 | 6.833 | 0.293 |
| NV0221 | GRANITE RANGE NV | 2011 | 40 | 0.760 | 0.737 | 4.162 | 85 | 7.083 | 0.353 |
| NV0222 | CALICO MOUNTAINS NV | 2005 | 22 | 0.685 | 0.733 | 4.043 | 70 | 5.833 | 0.129 |
| NV0222 | CALICO MOUNTAINS NV | 2010 | 42 | 0.768 | 0.747 | 4.272 | 90 | 7.500 | 0.356 |
| NV0222 | CALICO MOUNTAINS NV | 2011 | 40 | 0.748 | 0.723 | 4.009 | 83 | 6.917 | 0.337 |
| NV0226 | WARM SPRINGS CANYON NV | 2001 | 26 | 0.792 | 0.769 | 4.662 | 86 | 7.167 | 0.233 |
| NV0226 | WARM SPRINGS CANYON NV | 2012 | 27 | 0.701 | 0.742 | 4.361 | 85 | 7.083 | 0.235 |
| NV0234 | BLACK ROCK EAST NV | 2005 | 24 | 0.712 | 0.695 | 3.675 | 81 | 6.750 | 0.358 |
| NV0234 | BLACK ROCK EAST NV | 2010 | 28 | 0.690 | 0.719 | 3.814 | 76 | 6.333 | 0.237 |
| NV0234 | BLACK ROCK EAST NV | 2011 | 31 | 0.710 | 0.753 | 4.266 | 84 | 7.000 | 0.274 |
| NV0234 | BLACK ROCK WEST NV | 2005 | 25 | 0.700 | 0.708 | 3.795 | 80 | 6.667 | 0.313 |
| NV0234 | BLACK ROCK WEST NV | 2010 | 28 | 0.708 | 0.707 | 3.732 | 75 | 6.250 | 0.227 |
| NV0234 | BLACK ROCK WEST NV | 2011 | 19 | 0.675 | 0.654 | 3.226 | 68 | 5.667 | 0.265 |
| NV0507 | WHEELER PASS NV | 2002 | 23 | 0.777 | 0.700 | 3.593 | 58 | 4.833 | 0.103 |
| NV0507 | WHEELER PASS NV | 2007 | 26 | 0.756 | 0.763 | 4.636 | 85 | 7.083 | 0.259 |
| NV0612 | FISH CREEK NV | 2005 | 23 | 0.790 | 0.758 | 4.529 | 84 | 7.000 | 0.298 |
| NV0612 | FISH CREEK NV | 2014 | 183 | 0.781 | 0.782 | 5.287 | 98 | 8.167 | 0.327 |
| NV0618 | STONE CABIN NV | 2007 | 50 | 0.763 | 0.775 | 4.712 | 92 | 7.667 | 0.337 |
| NV0618 | STONE CABIN NV | 2012 | 46 | 0.792 | 0.783 | 4.827 | 91 | 7.583 | 0.253 |
| NV0619 | REVEILLE NV | 2010 | 51 | 0.753 | 0.715 | 3.777 | 78 | 6.500 | 0.295 |
| NV0619 | REVELLIE NV | 2014 | 44 | 0.723 | 0.718 | 3.842 | 78 | 6.500 | 0.282 |
| NV0619 | REVEILLE NV | 2017 | 68 | 0.734 | 0.734 | 4.103 | 86 | 7.167 | 0.326 |
| NV0620 | SAULSBURY NV | 2007 | 25 | 0.773 | 0.768 | 4.654 | 82 | 6.833 | 0.183 |
| NV0620 | SAULSBURY NV | 2012 | 44 | 0.731 | 0.769 | 4.559 | 85 | 7.083 | 0.259 |
| NV0621 | PAYMASTER NV | 2006 | 26 | 0.760 | 0.710 | 3.779 | 68 | 5.667 | 0.176 |
| NV0621 | PAYMASTER NV | 2010 | 49 | 0.748 | 0.702 | 3.742 | 70 | 5.833 | 0.186 |
| OR0002 | BEATYS BUTTE OR | 2002 | 42 | 0.676 | 0.770 | 4.585 | 92 | 7.667 | 0.304 |
| OR0002 | BEATYS BUTTE OR | 2010 | 32 | 0.747 | 0.763 | 4.550 | 80 | 6.667 | 0.213 |
| OR0002 | BEATYS BUTTE OR | 2018 | 34 | 0.772 | 0.772 | 4.558 | 89 | 7.417 | 0.326 |
| OR0003 | SOUTH STEENS OR | 2004 | 41 | 0.784 | 0.761 | 4.466 | 89 | 7.417 | 0.315 |
| OR0003 | SOUTH STEENS OR | 2010 | 31 | 0.758 | 0.741 | 4.137 | 83 | 6.917 | 0.301 |
| OR0003 | SOUTH STEENS OR | 2016 | 50 | 0.758 | 0.742 | 4.196 | 83 | 6.917 | 0.313 |
| OR0003 | SOUTH STEENS OR | 2018 | 30 | 0.731 | 0.724 | 3.768 | 79 | 6.583 | 0.329 |
| OR0004 | SHEEPSHEAD OR | 2002 | 60 | 0.704 | 0.781 | 4.827 | 87 | 7.250 | 0.195 |
| OR0004 | SHEEPSHEAD OR | 2011 | 48 | 0.790 | 0.787 | 4.870 | 87 | 7.250 | 0.264 |
| OR0007 | WARM SPRINGS OR | 2001 | 56 | 0.803 | 0.789 | 5.014 | 93 | 7.750 | 0.290 |
| OR0007 | WARM SPRINGS OR | 2010 | 83 | 0.766 | 0.778 | 4.607 | 96 | 8.000 | 0.344 |
| OR0007 | WARM SPRINGS OR | 2019 | 50 | 0.770 | 0.764 | 4.420 | 89 | 7.417 | 0.337 |
| OR0009 | RIDDLE MTN OR | 2003 | 14 | 0.762 | 0.698 | 3.612 | 63 | 5.250 | 0.143 |
| OR0009 | RIDDLE MTN OR | 2009 | 13 | 0.724 | 0.677 | 3.407 | 65 | 5.417 | 0.215 |
| OR0009 | RIDDLE MTN OR | 2011 | 20 | 0.717 | 0.657 | 3.221 | 66 | 5.500 | 0.318 |
| OR0009 | RIDDLE MTN OR | 2015 | 29 | 0.733 | 0.670 | 3.369 | 65 | 5.417 | 0.169 |
| OR0010 | KIGER HERD OR | 2009 | 12 | 0.729 | 0.688 | 3.391 | 63 | 5.250 | 0.238 |
| OR0010 | KIGER HERD OR | 2011 | 40 | 0.694 | 0.694 | 3.530 | 69 | 5.750 | 0.246 |
| OR0010 | KIGER HERD OR | 2015 | 56 | 0.722 | 0.707 | 3.710 | 80 | 6.667 | 0.300 |
| OR0011 | HOG CREEK OR | 2003 | 14 | 0.815 | 0.739 | 4.048 | 72 | 6.000 | 0.194 |
| OR0011 | HOG CREEK OR | 2018 | 33 | 0.798 | 0.745 | 4.340 | 75 | 6.250 | 0.147 |
| OR0013 | COLD SPRINGS OR | 2005 | 34 | 0.790 | 0.777 | 4.648 | 81 | 6.750 | 0.259 |
| OR0013 | COLD SPRINGS OR | 2010 | 24 | 0.806 | 0.754 | 4.214 | 72 | 6.000 | 0.222 |
| OR0013 | COLD SPRINGS OR | 2018 | 41 | 0.748 | 0.767 | 4.461 | 81 | 6.750 | 0.296 |
| OR0014 | COYOTE LAKE OR | 2001 | 73 | 0.746 | 0.743 | 4.035 | 86 | 7.167 | 0.291 |
| OR0014 | COYOTE LAKE OR | 2011 | 50 | 0.792 | 0.766 | 4.578 | 88 | 7.333 | 0.250 |
| OR0015 | JACKIES BUTTE OR | 2000 | 31 | 0.665 | 0.709 | 3.834 | 83 | 6.917 | 0.301 |
| OR0015 | JACKIES BUTTE OR | 2011 | 40 | 0.750 | 0.742 | 4.129 | 86 | 7.167 | 0.314 |
| OR0019 | MURDERERS CREEK OR | 2001 | 17 | 0.620 | 0.661 | 3.212 | 62 | 5.167 | 0.161 |
| OR0019 | MURDERERS CREEK OR | 2009 | 71 | 0.696 | 0.707 | 3.566 | 75 | 6.250 | 0.267 |
| OR0019 | MURDERERS CREEK OR | 2012 | 51 | 0.696 | 0.689 | 3.342 | 68 | 5.667 | 0.279 |
| OR0019 | MURDERERS CREEK OR | 2019 | 50 | 0.717 | 0.758 | 4.426 | 78 | 6.500 | 0.244 |
| UT0241 | CEDAR MOUNTAIN UT | 2002 | 20 | 0.742 | 0.726 | 4.067 | 73 | 6.083 | 0.260 |
| UT0241 | CEDAR MOUNTAIN UT | 2017 | 97 | 0.740 | 0.739 | 4.451 | 94 | 7.833 | 0.330 |
| UT0441 | BLAWN WASH UT | 2001 | 29 | 0.645 | 0.621 | 3.093 | 69 | 5.750 | 0.362 |
| UT0441 | BLAWN WASH UT | 2018 | 25 | 0.683 | 0.724 | 3.786 | 70 | 5.833 | 0.214 |
| UT0447 | NORTH HILLS UT | 2002 | 28 | 0.807 | 0.710 | 3.656 | 72 | 6.000 | 0.278 |
| UT0447 | NORTH HILLS UT | 2020 | 47 | 0.691 | 0.705 | 3.693 | 70 | 5.833 | 0.214 |
| UT0448 | SULPHUR N | 2006 | 56 | 0.740 | 0.736 | 3.933 | 80 | 6.667 | 0.338 |
| UT0448 | SULPHUR N | 2009 | 53 | 0.682 | 0.732 | 3.964 | 76 | 6.333 | 0.303 |
| UT0448 | SULPHUR S | 2006 | 12 | 0.625 | 0.679 | 3.613 | 65 | 5.417 | 0.169 |
| UT0448 | SULPHUR S | 2009 | 41 | 0.679 | 0.715 | 3.723 | 70 | 5.833 | 0.271 |
| UT0641 | CEDAR RIDGE UT | 2002 | 30 | 0.667 | 0.625 | 2.838 | 53 | 4.417 | 0.170 |
| UT0641 | CEDAR RIDGE UT | 2006 | 14 | 0.702 | 0.655 | 3.010 | 51 | 4.250 | 0.157 |
| UT0641 | RANGE CREEK UT | 2006 | 26 | 0.663 | 0.707 | 3.633 | 63 | 5.250 | 0.159 |
| UT0641 | RANGE CREEK UT | 2018 | 50 | 0.682 | 0.652 | 3.005 | 65 | 5.417 | 0.246 |
| WY0002 | DIVIDE BASIN WY | 2004 | 50 | 0.797 | 0.793 | 4.959 | 89 | 7.417 | 0.292 |
| WY0002 | DIVIDE BASIN WY | 2011 | 60 | 0.785 | 0.787 | 4.929 | 93 | 7.750 | 0.333 |
| WY0009 | ADOBE TOWN WY | 2003 | 40 | 0.773 | 0.771 | 4.681 | 90 | 7.500 | 0.278 |
| WY0009 | ADOBE TOWN WY | 2010 | 103 | 0.776 | 0.776 | 4.748 | 93 | 7.750 | 0.280 |
| WY0009 | ADOBE TOWN WY | 2017 | 25 | 0.790 | 0.778 | 4.836 | 76 | 6.333 | 0.132 |
| WY0027 | MUSKRAT BASIN WY | 2004 | 27 | 0.731 | 0.727 | 3.851 | 72 | 6.000 | 0.222 |
| WY0027 | MUSKRAT BASIN WY | 2013 | 77 | 0.759 | 0.765 | 4.440 | 81 | 6.750 | 0.222 |
| WY0028 | DISHPAN BUTTE WY | 2004 | 30 | 0.768 | 0.743 | 4.099 | 76 | 6.333 | 0.211 |
| WY0028 | DISHPAN BUTTE WY | 2013 | 30 | 0.756 | 0.726 | 4.002 | 70 | 5.833 | 0.200 |
| WY0029 | CONANT CREEK WY | 2004 | 22 | 0.680 | 0.656 | 3.394 | 66 | 5.500 | 0.288 |
| WY0029 | CONANT CREEK WY | 2013 | 28 | 0.714 | 0.688 | 3.590 | 72 | 6.000 | 0.208 |
| WY0031 | ANTELOPE HILLS WY | 2004 | 25 | 0.747 | 0.733 | 3.914 | 76 | 6.333 | 0.237 |
| WY0031 | ANTELOPE HILLS WY | 2012 | 34 | 0.767 | 0.758 | 4.343 | 87 | 7.250 | 0.333 |
| WY0035 | LOST CREEK WY | 2006 | 48 | 0.767 | 0.791 | 5.015 | 84 | 7.000 | 0.190 |
| WY0035 | LOST CREEK WY | 2009 | 30 | 0.775 | 0.788 | 4.899 | 80 | 6.667 | 0.200 |
| WY0039 | LITTLE COLORADO WY | 2007 | 36 | 0.801 | 0.762 | 4.363 | 79 | 6.583 | 0.190 |
| WY0039 | LITTLE COLORADE WY | 2011 | 45 | 0.761 | 0.768 | 4.541 | 80 | 6.667 | 0.188 |

Various measures of genetic distance and genic differentiation among the populations were examined and genetic distance was compared to geographic distance based upon GPS coordinates for each herd (supplied by the BLM). In general, there was no strong correlation of genetic distance with geographic distance but those herds that are geographically closest to each other average lower genetic distances. We tested for a correlation of geographic distance to genetic distance (as measured by Nei’s Da distance) with the Mantel test. The measures did show a highly statistically significant correlation (p< 0.0001) however, the correlation was low (r=0.181) which means the association explained less than 3.3 percent of the total variability. Figure 1 shows the relationship of most of the herds to each other geographically. Visualization of genetic relationships was done by construction of Restricted Maximum Likelihood trees and Factorial Correspondence analysis (Figures 2, 4 and 5). There is some tendency for herds from within the same state to cluster together (for example herds from CO and ID in Figure 2), however in many cases, although the herds in a cluster are from the same state they are not really in close geographic proximity. There are several clusters that are predominately made up of Herds from Nevada but to some extent this is due to the large number of herds from that state and not close relationship. Herds from California were scattered throughout the tree. In the FCA analyses, there are some clear outliers which are mostly herds with variability levels outside those near the mean values. The outlier herds tended to be those with low sample size which also tended to be those with low AML values. Figure 5 shows the FCA plot with some of the outlier herds removed to give a clearer view of relationships.

**Figure 2.**
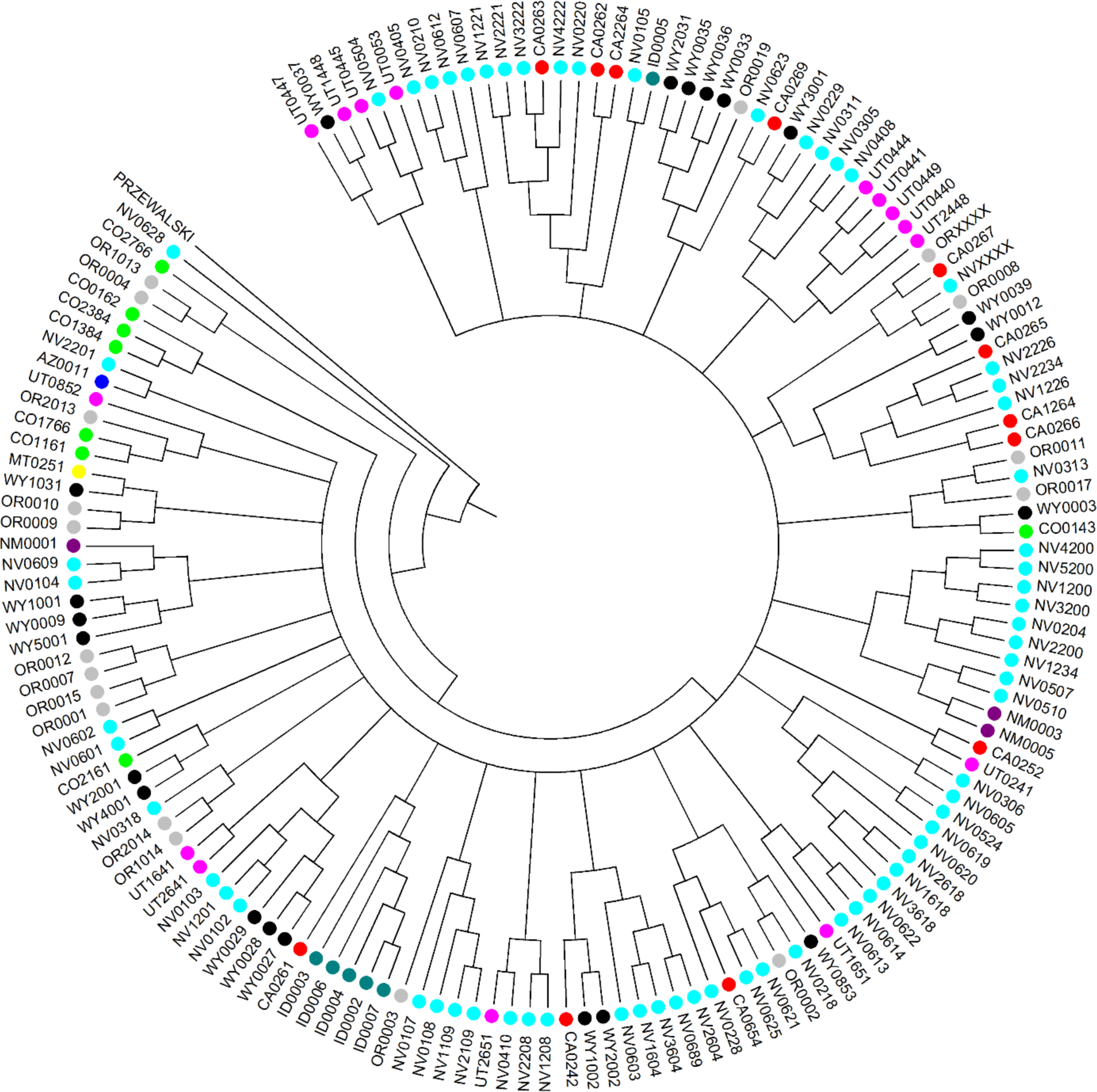
Restricted Maximum Likelihood dendrogram based upon Chord Distance of feral herds (152) color coded by state.

**Figure 3.**
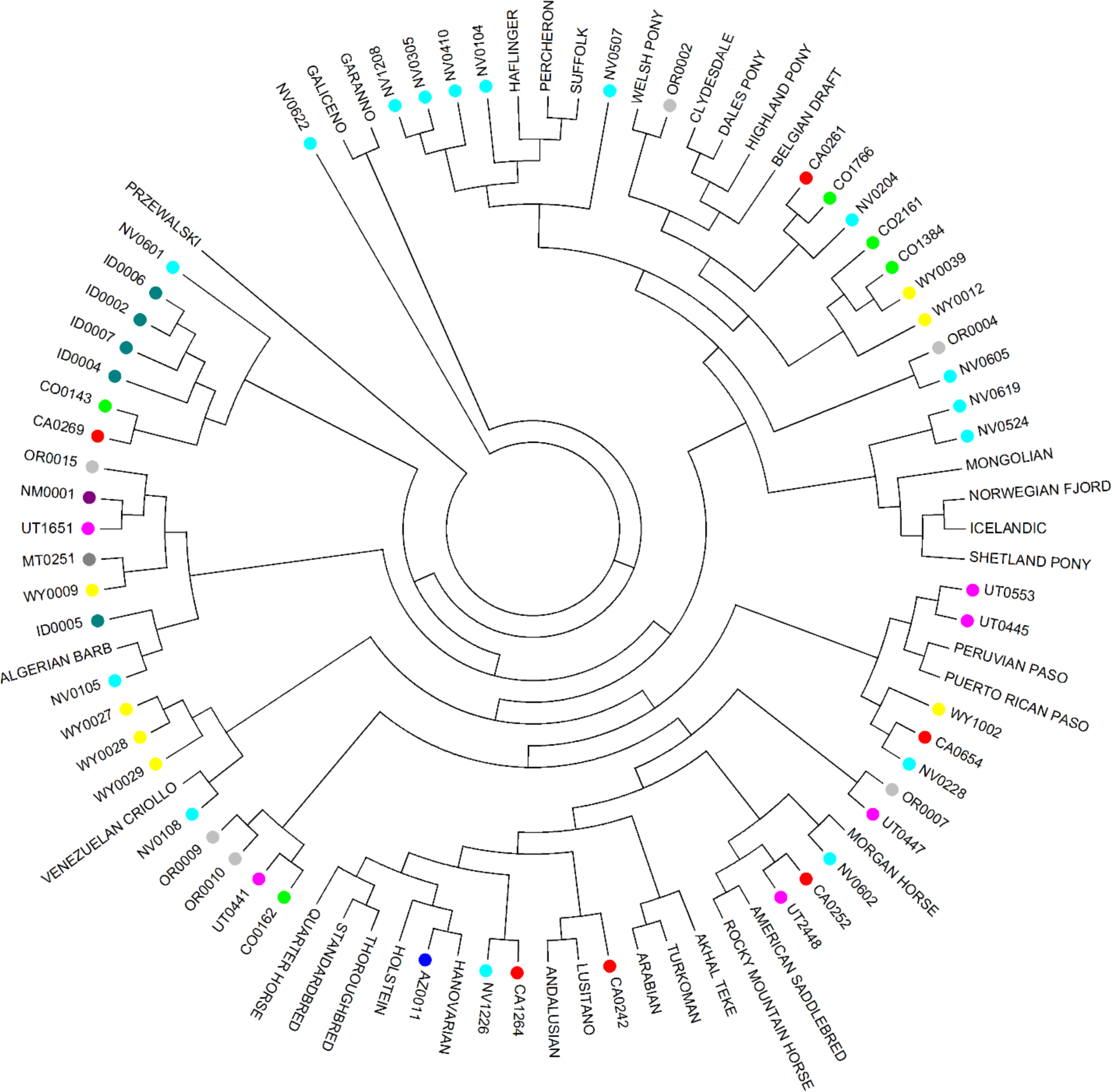
RML dendrogram from Chord distance of 55 selected feral herds and 31 domestic breeds plus Przewalski Horse as an out-group.

**Figure 4.**
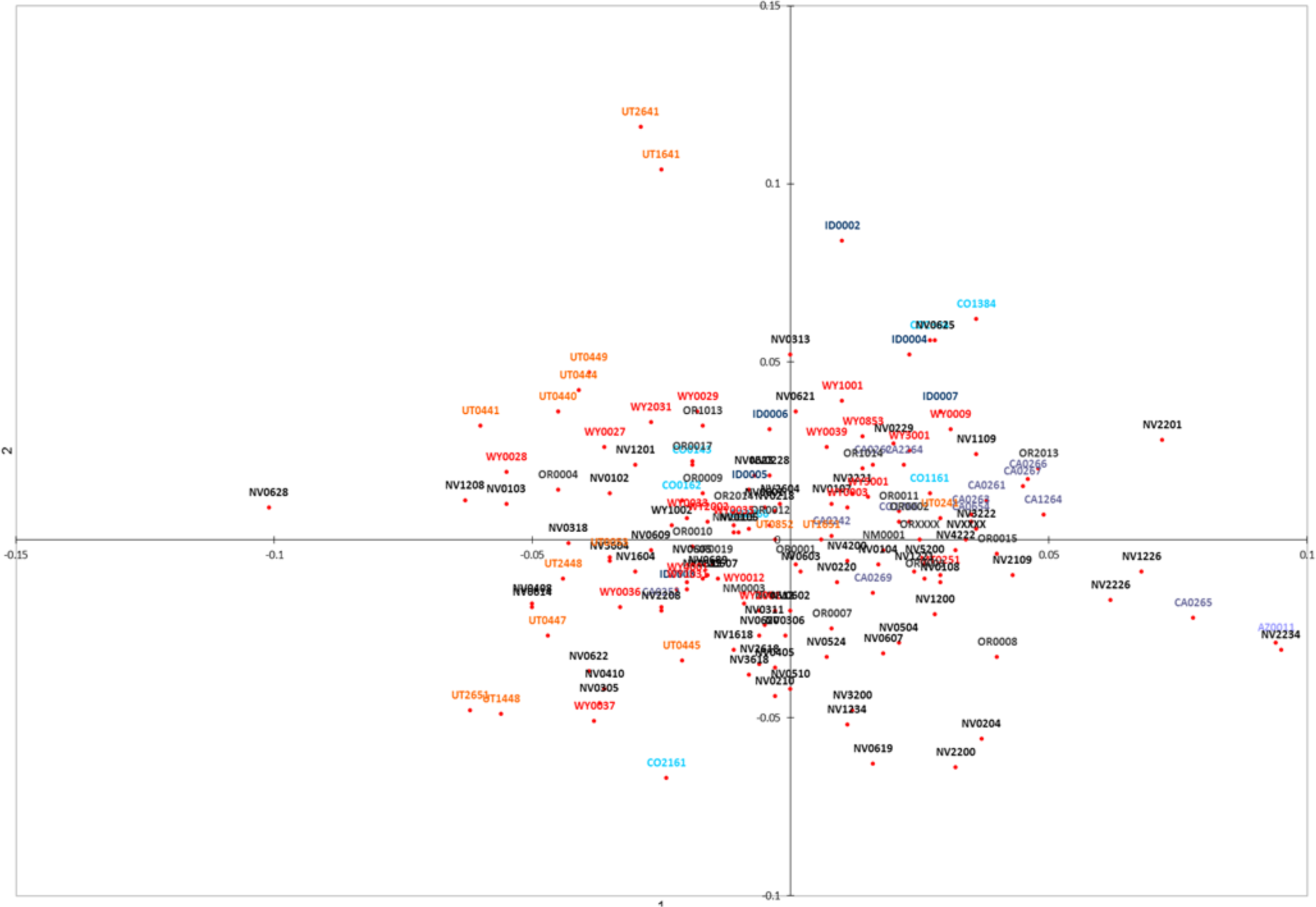
PCoA using the same 152 populations in Figure 2. Only axes 1 and 2 are plotted.

**Figure 5.**
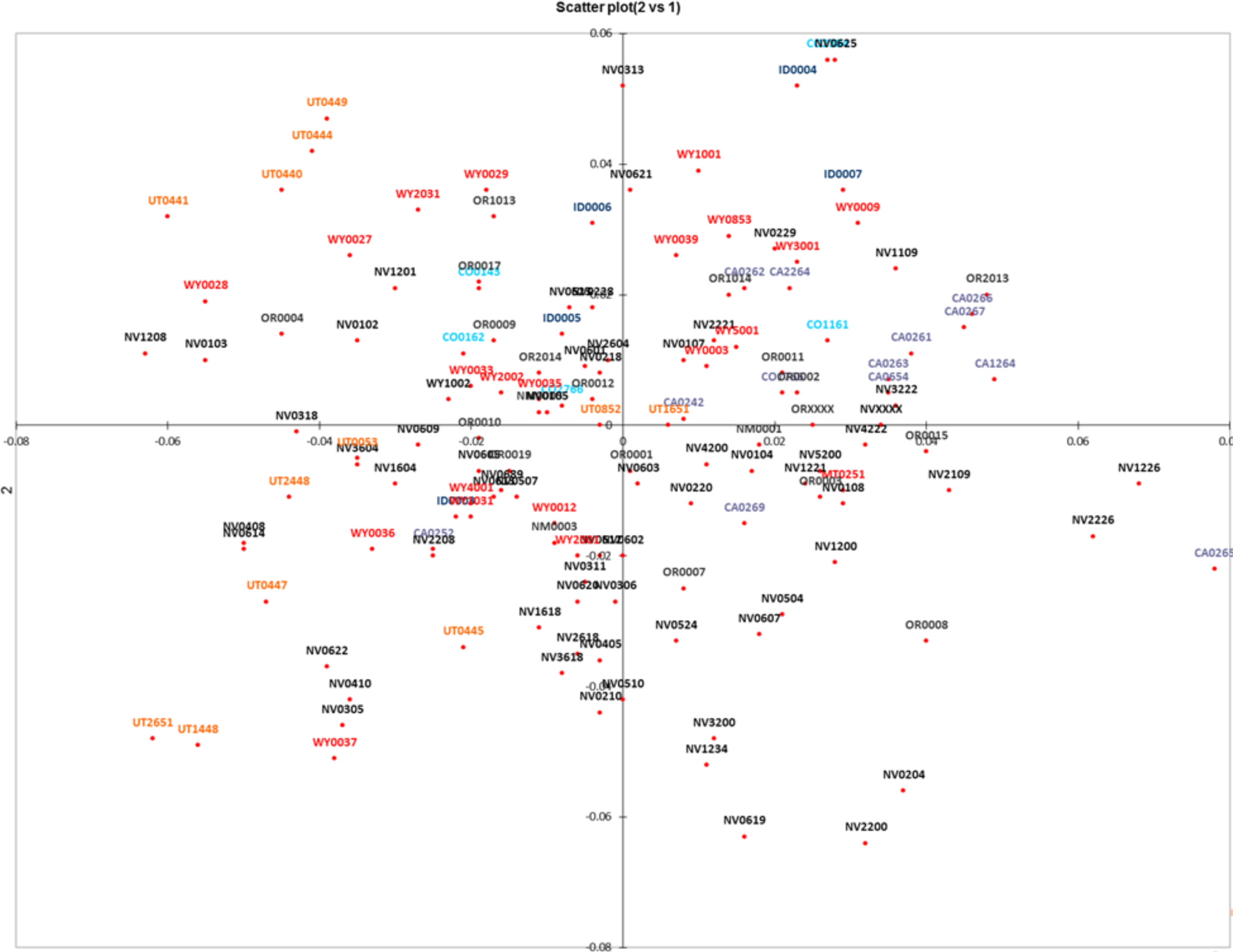
PCoA (axes 1 and 2) based upon removal of outlier herds UT2641, UT1641, CO2161, CO1384, NV0628, AZ0011, NV2234, NV2201 and ID0002 but otherwise using the same populations as Fig. 4.

A heat diagram of the Fst values is shown in Figure 6. What this diagram shows is the relative degree of differentiation of each herd compared to all others. By following the row and/or column for each herd the degree of differentiation can be determined by the color of the line. The darker the line the higher the differentiation while the lighter the line the lower. Almost all herds with very dark lines had a small sample size thus differentiation was based largely on sample dependent variability measures. The first herd on the chart is the AZ0011-Cibola Trigo herd which had an acceptable sample size but the lowest levels of variation of any of the herds tested. The other herd that stands out is NV0628-Gold Mountain and this herd also has quite low variation levels.

**Figure 6.**
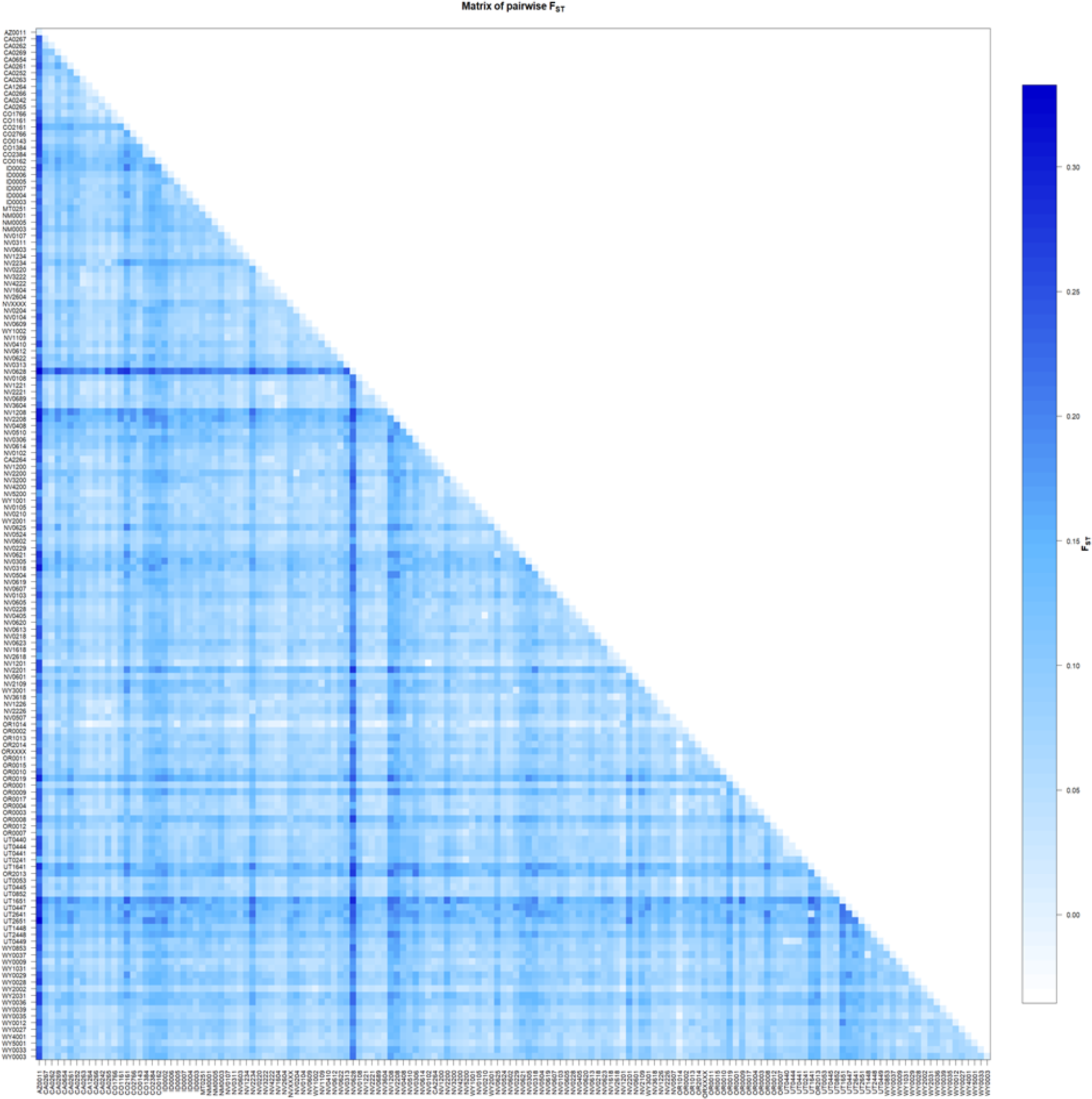
Pairwise Fst matrix.

STRUCTURE and FLOCK analyses were performed to test for any genetic pattern or substructure among all populations. STUCTURE did not produce any visually clearly identifiable population structure at K=2 (Figure 7). Twelve sample populations were assigned with more than 90% to the ancestral population cluster “orange” in Figure 7: CIBOLO TRIGO AZ0011 (sample population number 1), JICARILLA NM0003 (31), DESATOYA MTN NV0606 (44), LITTLE OWHYEE TUS NV0200 (68), REVEILLE NV0619 (84), ROCKY HILLS NV0605 (87), STONE CABIN S NV0618(94), STONE CABIN N NV0618(95), STONE CABIN 2007 NV0618 (101), NORTH HILLS UT0447 (138), SULPHUR N UT0448 (141), and COOPER CREEK WY00370037 (145). With the exception of sample populations 68, 87 and 138, which showed increasing admixture, all other populations remained clustered with each other at higher K values (e.g. dark violet at K=8). At K=2, twelve populations were assigned with more than 90% to the ancestral population cluster “blue” in Figure 7: BITNER CA0267 (sample population number 3), CATNIP NVXXXX (42), NORTH STILLWATER NV0229 (79), SILVER PEAK NV0623 (93), STILLWATER NV (100), LIGGETT TABLE OR0037 (116), BIBLE SPRINGS UT0440 (127), 4 MILE UT0444 (128), CEDAR RIDGE TRAP UT0461 (131), COLD SPRINGS TRAP OR0013 (132), COLD SPRINGS OR0013 (133), and RANGE CREEK UT0641 (139). All of these showed increasing admixture with increasing K with the exception of COLD SPRINGS TRAP OR and CEDAR RIDGE UT (sample populations 131 and 133, “red” ancestral population at K=8) and LIGGETT TABLE OR (sample population 116, “light green” ancestral population at K=8). At K=8, one set of populations emerges with the “dark green” ancestral population of larger than 75%: FOX HOG CA0263 (sample population 9), CALICO MTNS NV0222 (38) and CALICO NV0222 (39). These results clearly do not show any pattern of population genetic clustering nor of geographic relationships except for those herds which are essentially in the same localities (94, 95, 101 and 79, 101)

**Figure 7.**
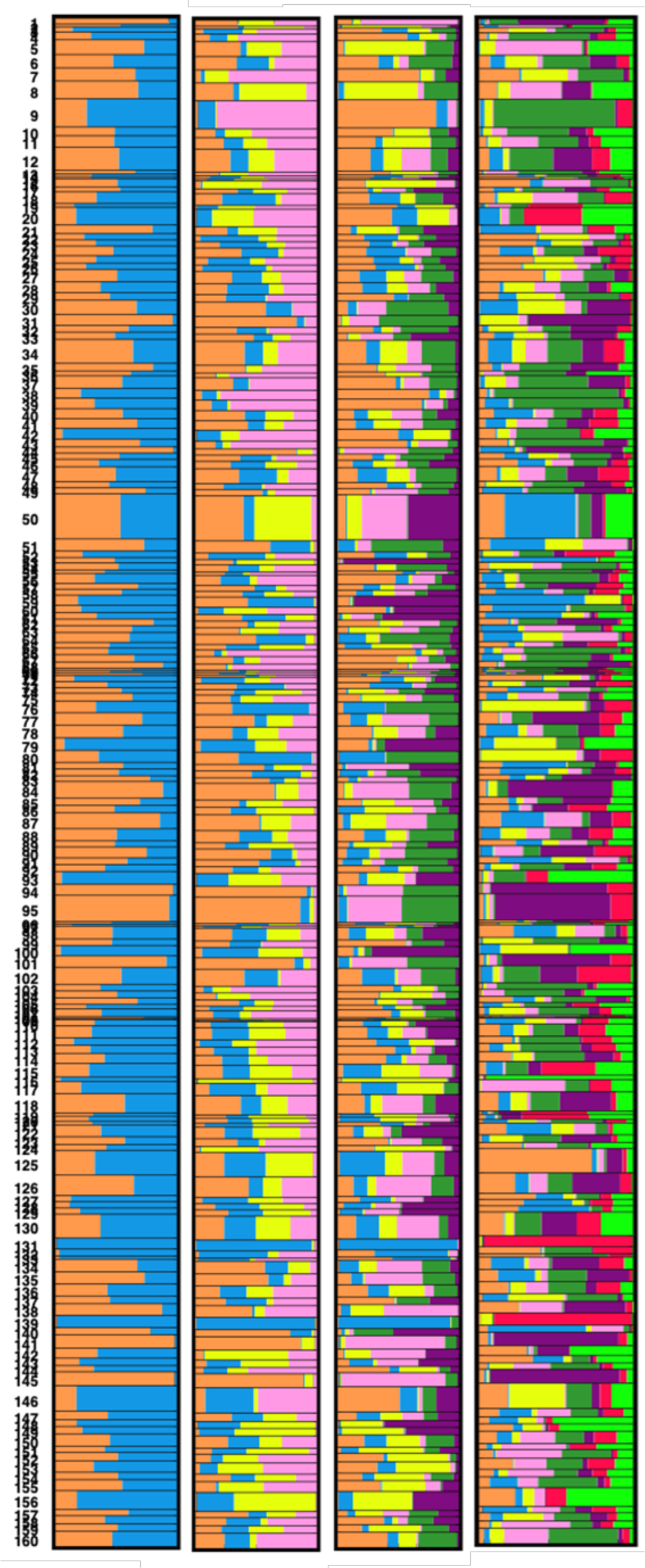
Structure analysis for K=2, K=4, K=6, and K=8, respectively (from left to right).

No FLOCK stopping condition was reached when all populations were evaluated at K=2. We removed all 97 sample populations with less than 80% of horses allocated to a single population cluster and repeated the analysis with the remaining 63 sample populations. The repeat produced again no stopping condition. Stepwise increasing K to K=6 did not result in any stopping condition. This indicates no geographic clustering.

Possible ancestry of the feral herds to domestic breeds was estimated by constructing a RML tree from 55 selected feral herds and 31 domestic horse breeds with a focus on those from the New World (Figure 3). Most herds appeared to be of widely mixed ancestry with no clear, specific breed ancestor. This result is consistent with the other genetic results (STRUCTURE, FLOCK and FCA) and the history of most of the areas where the feral herds are located. All of these herds have been individually analyzed in reports provided to the BLM at the time the herds were sampled. These analyses consistently indicated that the herds had mixed-breed ancestry and the majority showed closest similarity to breeds of North American origin. Only a handful of herds show evidence of Iberian ancestry and those that do tend to be more isolated than other herds. Herds that were generally considered to be Spanish in origin were MT0251, OR0010 and UT0448, but these herds were not clustered with Spanish breeds in Figure 3 which is largely due to a change in the make-up of the herds over time.

Modularity analysis results indicate that there is moderate modularity in the modeled genetic networks, meaning there is connection that correlates with the spatial estimates provided by longitude and latitude of sampling location. The density ratio, the ratio of links within to between modules, was 0.5668 which suggests that there are 1.76x more connections between modules than within modules. This is indicative of considerable gene flow between sampling locations. Both the non-spatial (modularity = 0.261) and spatial (modularity = 0.244) tests for modularity were significant (P < 0.001, respectively). These results indicate that there is greater movement of individuals among herd locations within the same module than among herds between modules, and using physical distance between sampling locations decreased our estimate of modularity, also indicating significant genetic connections between locations.

The general trend for feral herds on public lands has been a decrease in population size over time. This will inevitably lead to a long-term loss of genetic variability within individual HMAs. However, this loss of variation can be mitigated by a low level of exchange of individuals from geographically close herds. This process will tend to homogenize the herds but this would take many generations. Table 4 shows the possible number of migrants from herds that were geographically close to another herd (less than 100 km). In some cases, it appeared that there were a very large number of individuals that possibly originated from a nearby herd rather than the one they were captured in. However, this may simply be due to closer genetic similarity of geographically close herds rather that migration as well as possible subdivision within a herd. The relationship of the possible number of migrants within a herd and genetic variability measure was analyzed. All measures showed a statistically highly significant association. The results of the analysis are shown in Figure 8. Some consideration should be made to manage a few of the herds in such a way that they are preserved as is, as much as possible, for historical or, in the cases of the small number of herds with strong Spanish horse genetics, for breed type conservation.

**Figure 8.**
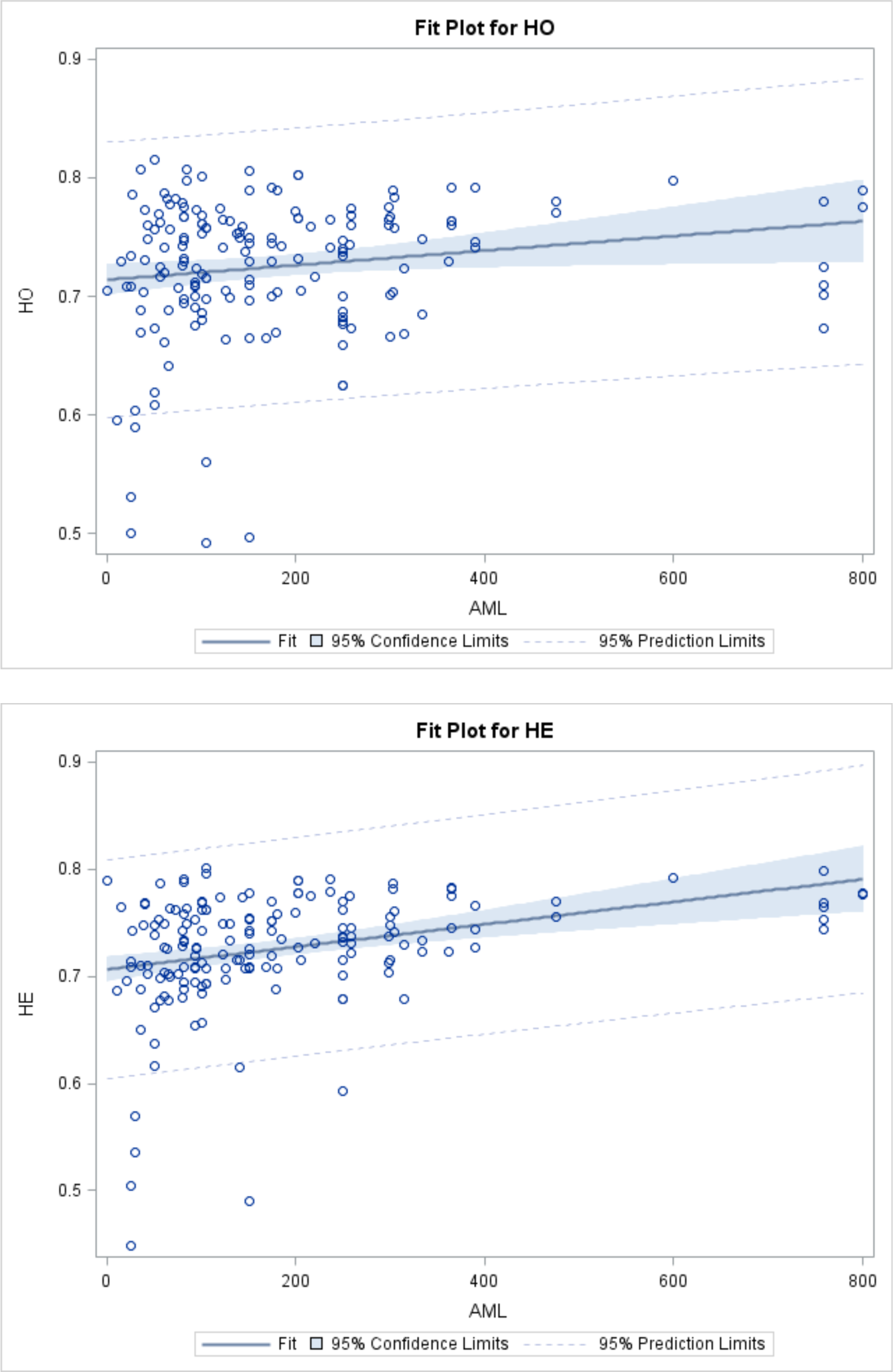
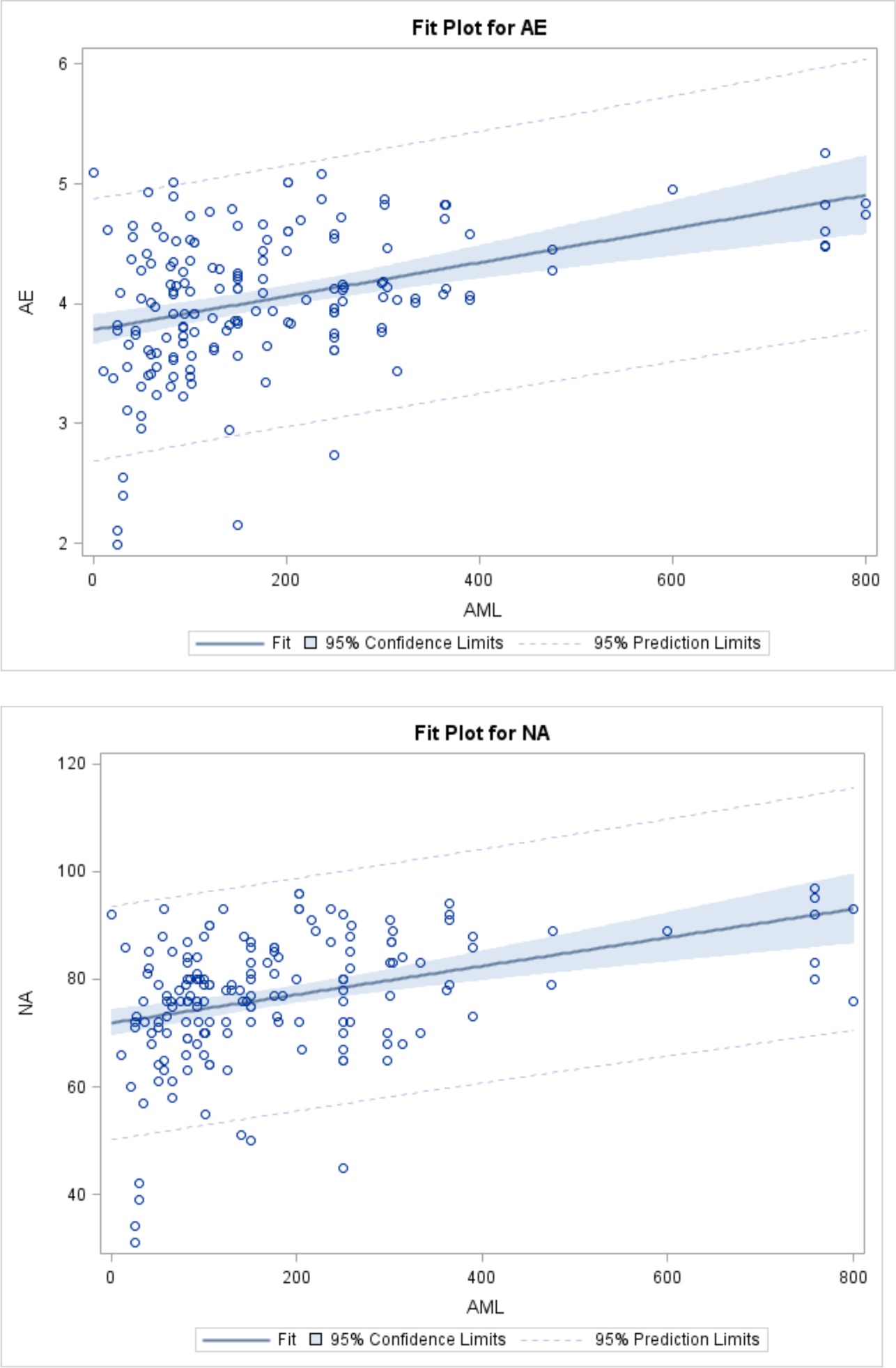
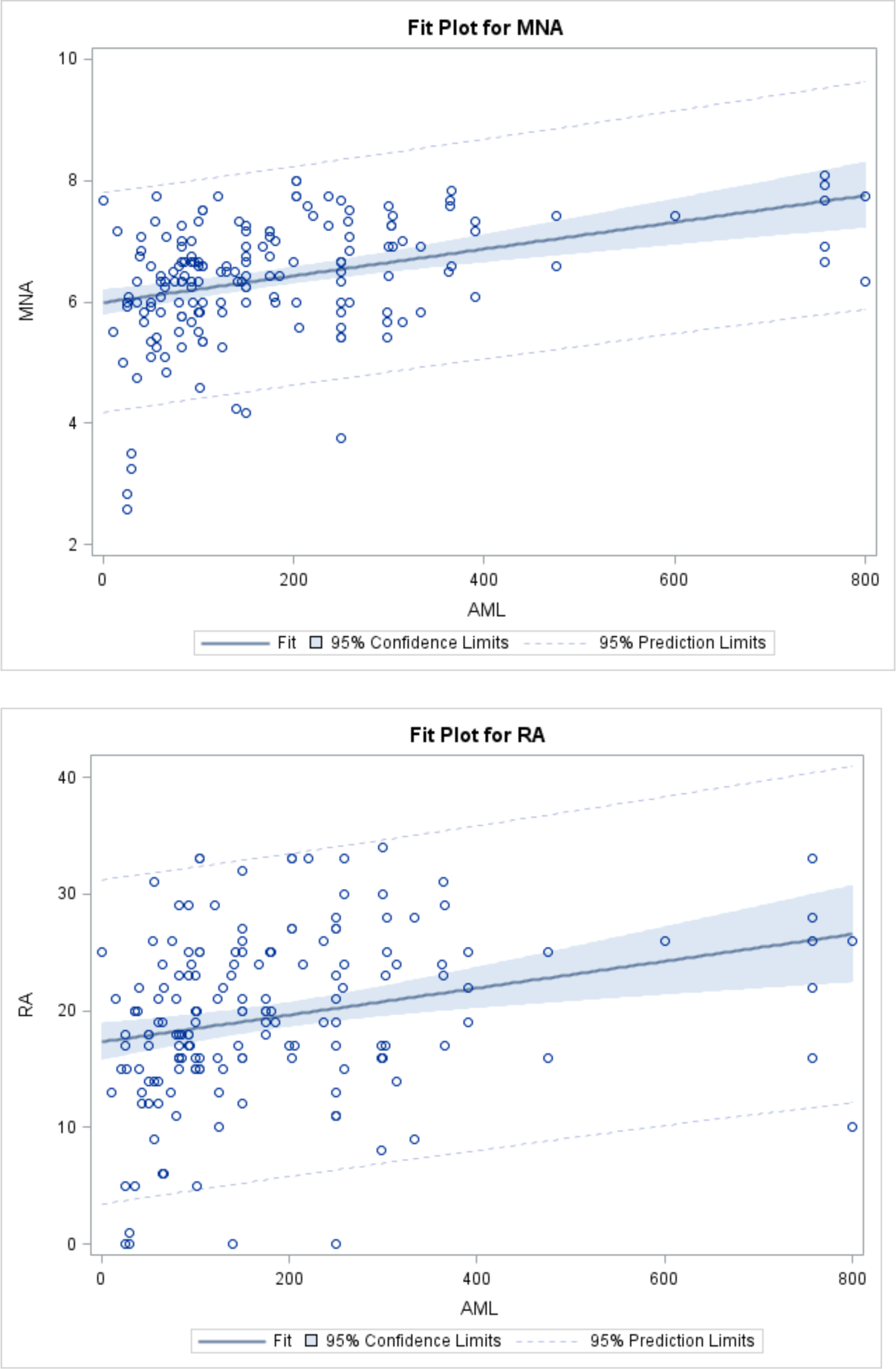
Plots of associations of different genetic measures with AML.

**Figure 9.**
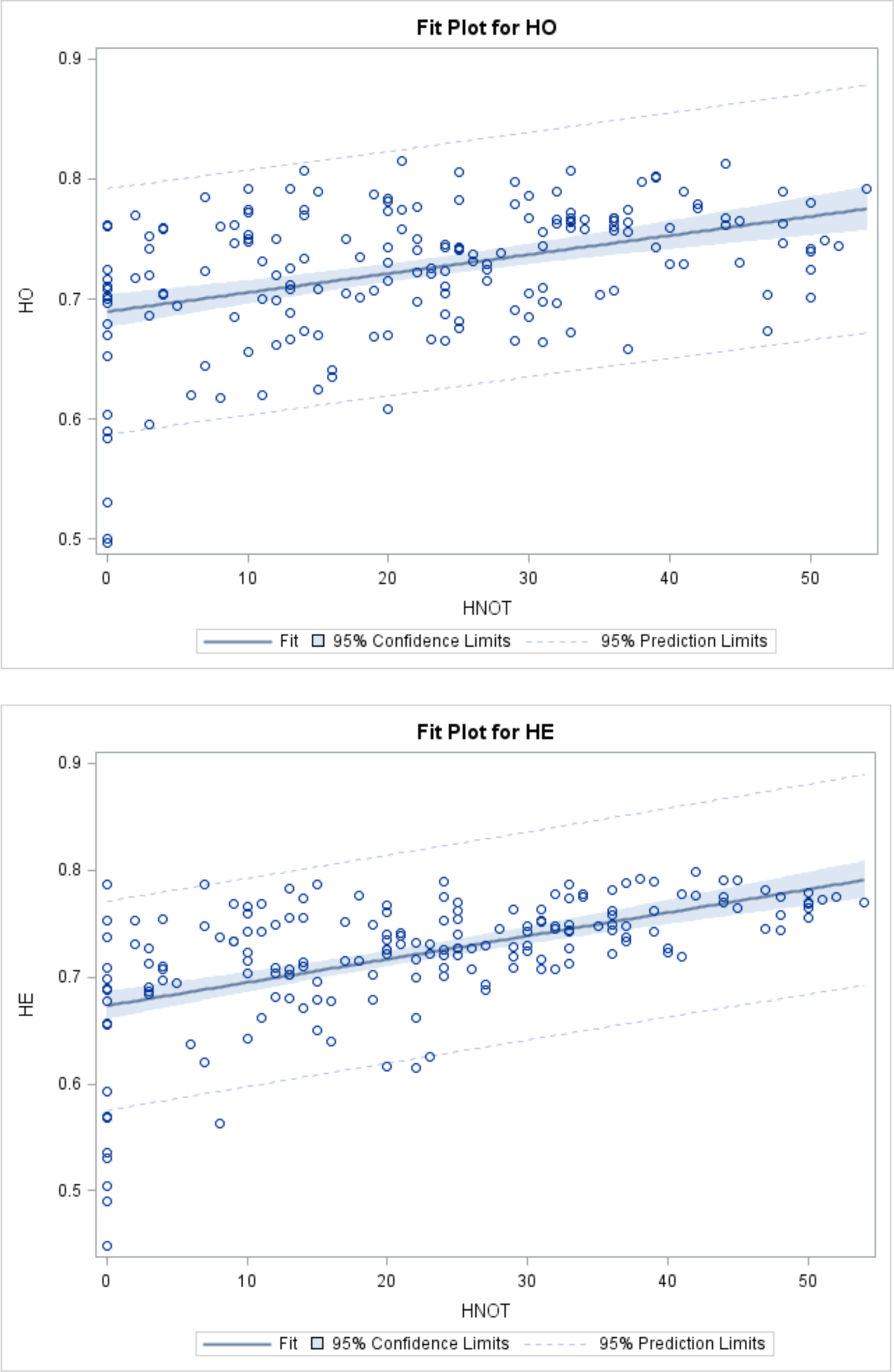
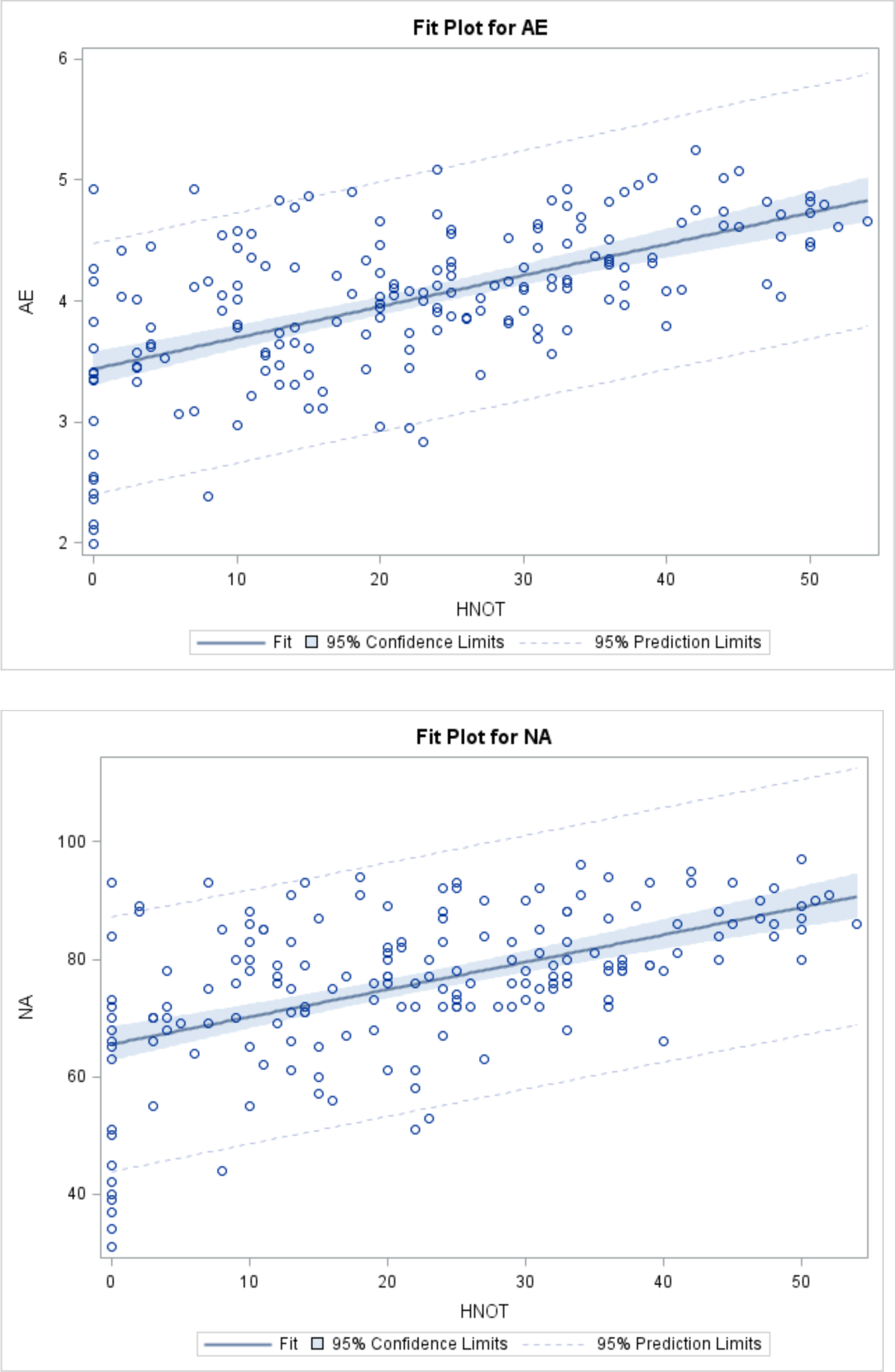
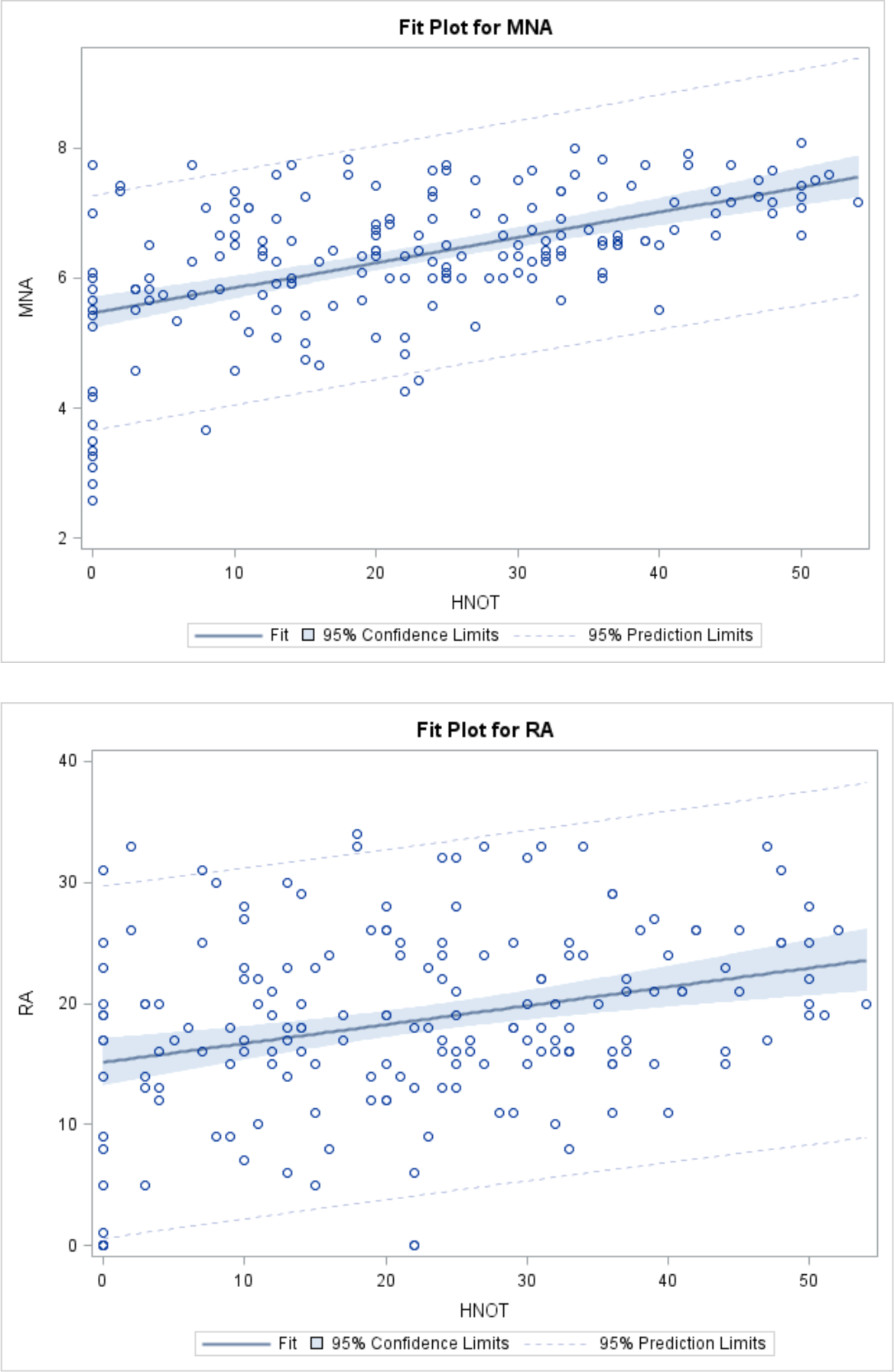
Plots of association of different genetic measures with percent of likely migrants.

**Table 4.**
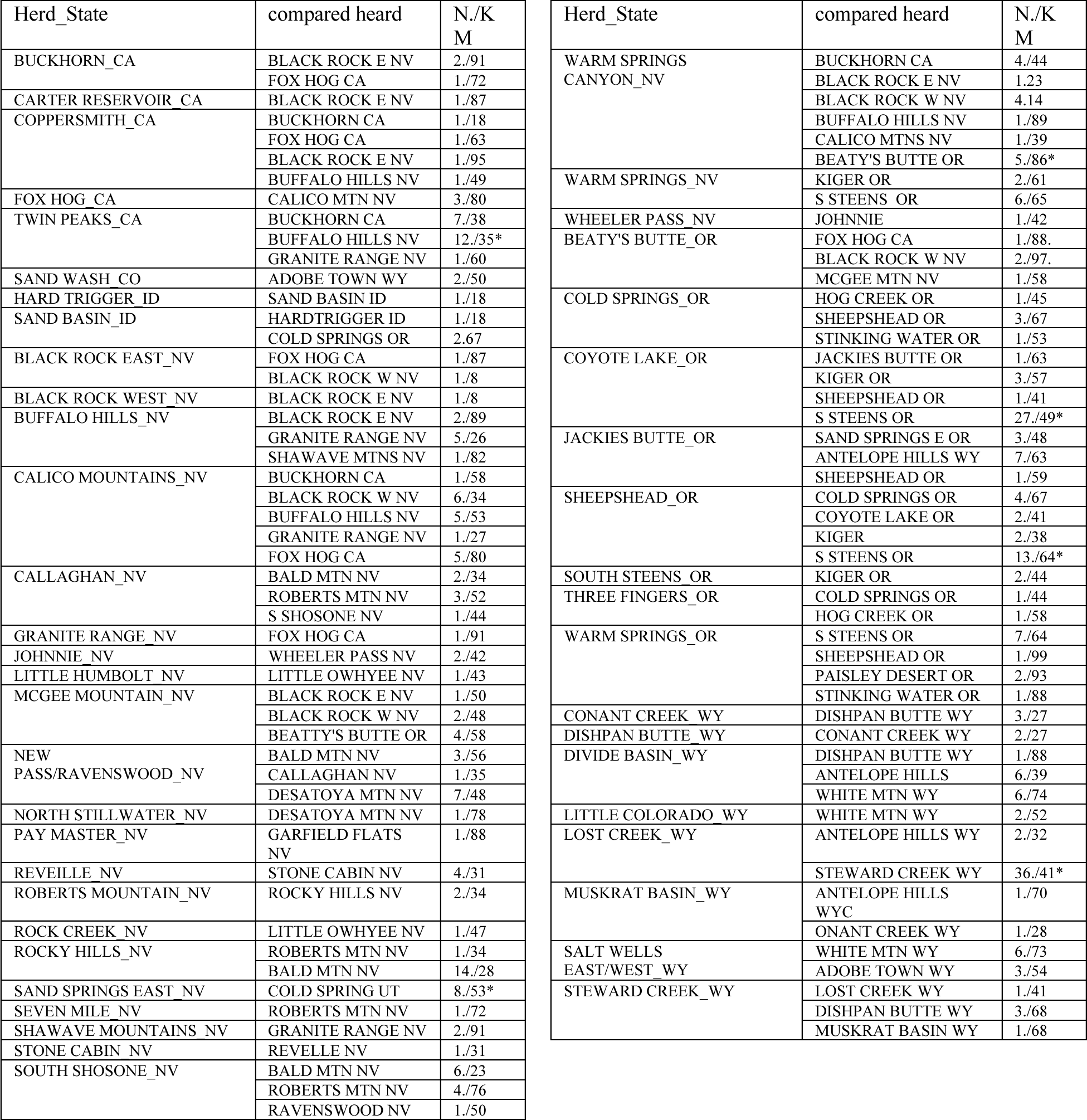
Estimates of the possible number of immigrants into a herd from nearby herds. N refers to the number of individuals and KM refers to the geographic distance in kilometers.

To examine possible loss of variation within herds we looked at variability measure in herds that were sampled more than one time. In general, there was no clear pattern to the differences in variability measures from time 1 to any other time period (Table 3). There are several possible reasons for this. One is that insufficient time has elapsed between two or more sampling periods. Changes in variability take place over the period of generations and are not likely to be observable over a period of 4 to 10 years. Another is that in some cases it is likely that there was introgression of individuals from an outside area which would increase variability from time 1 to time 2 or later. In some cases, this may have been done intentionally to increase the variation within a herd. Another possibility is simply differences in sample sizes in different years. In some cases, initial sample size was small and the second or later sample size was larger. The reverse also occurred.

This work shows that there is a great deal of genetic diversity within feral horse herds in the Western United States. The total diversity approaches that of domestic horse breeds. At least up to now, there has been no significant decrease in genetic variability over the past 20 years; however, population sizes of the herds is generally low so that loss of variation in these mostly closed herds is inevitable. Management practices applied by the responsible agencies can reduce the rate of loss of variation and practices such as intentional introductions of animals from outside herds are currently in use. Part of the basis for the current high levels of variation seen in many of the herds is that they have a mixed breed origin that depended upon what domestic breeds were in the area and became feral for whatever reason. There does not appear to be any large pattern of genetic structure across the total area where feral horses are found. There may be some small-scale structure within specific regions and this can be seen in the associations of herds within states seen in Figures 2 and 4. More work is needed to determine the scale of this structure and to understand the basis for it.

## Acknowledgments

I wish to thank all the people with the Bureau of Land Management that helped make this work possible, in particular Dr. Paul Griffin, Alan Shepard, Dean Bolstad, Joseph Stratton, and Bea Wade. A special thanks to Linda Coates-Markle for her friendship and support. This work has been supported by funds from the BLM since 1992 in various forms and current work under Federal Assistance contract SF-424, 2009 and 2011.

